# Electrostatic–hydrophobic cooperation in the mesoscale assembly of synapsin and synaptic vesicles

**DOI:** 10.64898/2026.11.12.766173

**Authors:** Shokoofeh Akbari, Oleg Shupliakov, Yousef Jamali

**Affiliations:** Department of Applied Mathematics, School of Mathematical Sciences, Tarbiat Modares University, Tehran, Iran; Department of Neuroscience, Karolinska Institutet, Stockholm, Sweden

## Abstract

Synapsin associates with synaptic vesicles. How these local contacts give rise to a connected vesicle assembly remains unclear. We used a three-dimensional Brownian agent model to vary electrostatic and hydrophobic interactions independently at fixed particle abundance and confinement. With electrostatics alone, the agents representing synapsin approached almost every vesicle early, when most vesicles still belonged to separate groups. With both interactions active, a larger connected assembly formed earlier, more of these agents were near several vesicles, and they moved more slowly. Over the full 84 ms, the assemblies continued to grow under either interaction alone. By the final observation window, the two conditions had similar vesicle participation despite differences in protein association and contact composition. The positive early interaction contrast decreased as the assemblies grew. These observations distinguish local synapsin association from collective vesicle connectivity and identify a structural feature that can be measured alongside condensate formation.

**Author summary:** Synaptic vesicles store the molecules that neurons use to communicate. The protein synapsin helps organize these vesicles into groups. We wanted to know how interactions at individual vesicles give rise to connections across a whole group. In our computer model, electrical interactions quickly brought the particles representing synapsin close to almost every vesicle. At that point most vesicles still belonged to separate groups. Adding hydrophobic attraction led to a larger connected group sooner and slowed the motion of these particles. We then followed the simulations for longer. The groups also grew when each interaction acted alone, and eventually similar fractions of vesicles belonged to the largest group in the two conditions, even though the protein contacts differed. These results show why it matters to measure which vesicles are linked across a group as well as how much protein is nearby. Measuring these properties alongside the formation of protein-rich condensates could help explain how synapsin organizes vesicles.

## 1 Introduction

A presynaptic terminal holds synaptic vesicles (SVs) in groups while allowing them to move and participate in neurotransmission. The protein-rich SV membrane offers many sites for local interactions [1]. Association at individual vesicles and connectivity across a vesicle cluster are different levels of organization. Understanding how local contacts link otherwise separate vesicle groups is therefore central to a physical description of clustering.

Experiments on synapsin have linked molecular interactions to vesicle organization. Early bilayer experiments identified both electrostatic association and hydrophobic engagement of the synapsin head domain [2]. Purified-vesicle experiments distinguished lipid binding from an ionicstrength and phosphorylation-sensitive protein-associated contribution [3]. Computational cage and cytoskeletal cross-linking models then examined how phosphorylation-dependent binding could redistribute vesicles [4]. Later, perturbation of an amphipathic lipid-packing sensor motif linked membrane curvature sensing to synapsin binding and vesicle clustering [5]. These experiments identify distinct molecular contributions to synapsin binding and vesicle organization.

Reconstitution experiments showed that synapsin assemblies capture vesicles and disperse upon kinase treatment [6]. Perturbing the disordered region of synapsin also affects vesicle clusters in a living lamprey synapse [7]. Single-molecule tracking linked vesicle sequestration to restricted synapsin motion [8]. In non-neuronal cells, co-expression of synapsin and synaptophysin generated SV-like clusters [9]. Endophilin A1 can also form condensates with synapsin and becomes incorporated into SV-like clusters [10]. Synaptic context also matters. Song and Augustine distinguished synapsin-dependent liquid–liquid phase separation (LLPS) at inhibitory synapses from tetramer-mediated cross-linking at excitatory synapses [11]. Reserve-pool tracking also implicated synapsin IIa tetramerization in vesicle organization [12]. These findings motivate separate measures of protein association, vesicle connectivity and particle motion.

Microscopy and a binding-particle model related assembly size and morphology to the synapsin to-lipid ratio and vesicle adhesion [13]. X-ray scattering and cluster simulations resolved noncom- pact SV–synapsin assemblies with close intervesicle contacts [14]. X-ray microrheology of synapsin assemblies with colloid-supported lipid bilayers found concentration-dependent motion and relax- ation [15].

Simulation approaches span several resolutions. Atomistic synapsin simulations coupled to experiments revealed an ATP-dependent conformational route to vesicle binding and oligomerization [16]. Residue-level models resolve charge and hydropathy along protein chains [17], while coarse-grained simulations of repetitive peptides show that the balance of electrostatic repulsion and hydrophobic attraction depends on sequence and solution conditions [18]. Agent-based diffusion models have examined how crowding and geometry affect particle motion [19]. At this resolution, we can track individual proteins and vesicles while following the formation of a network. Connectivity can change nonlinearly with the number and arrangement of contacts, as in random and geometric graphs [20, 21].

We use a three-dimensional Brownian agent model to examine how local synapsin–vesicle contacts develop into mesoscale connectivity. We vary electrostatic and hydrophobic interactions independently and hold particle abundance, confinement and dispersion attraction fixed. The contact networks distinguish vesicle coverage from connectivity across the assembly. We also compare short-lag synapsin mobility with published tracking data.

## 2 Results

We denote simulated synapsin agents by Syn. We compared four conditions with electrostatic (E) and hydrophobic (H) interactions switched on (1) or off (0). Each condition had three runs, one per seed block, followed for 84 ms. Structural summaries use the final 180 frames (58.94–84 ms). The same trajectories through 28 ms provide an earlier comparison.

### 2.1 Combined interactions produce an earlier and larger connected assembly

With neither E nor H active, the late mean all-agent largest-component fraction was 0.0368. It reached 0.3440 with H alone, 0.5025 with E alone and 0.9200 with both interactions (Table 1 and Fig 1). The combined-interaction condition had the largest component in every seed at both observation horizons.

**Fig 1.**
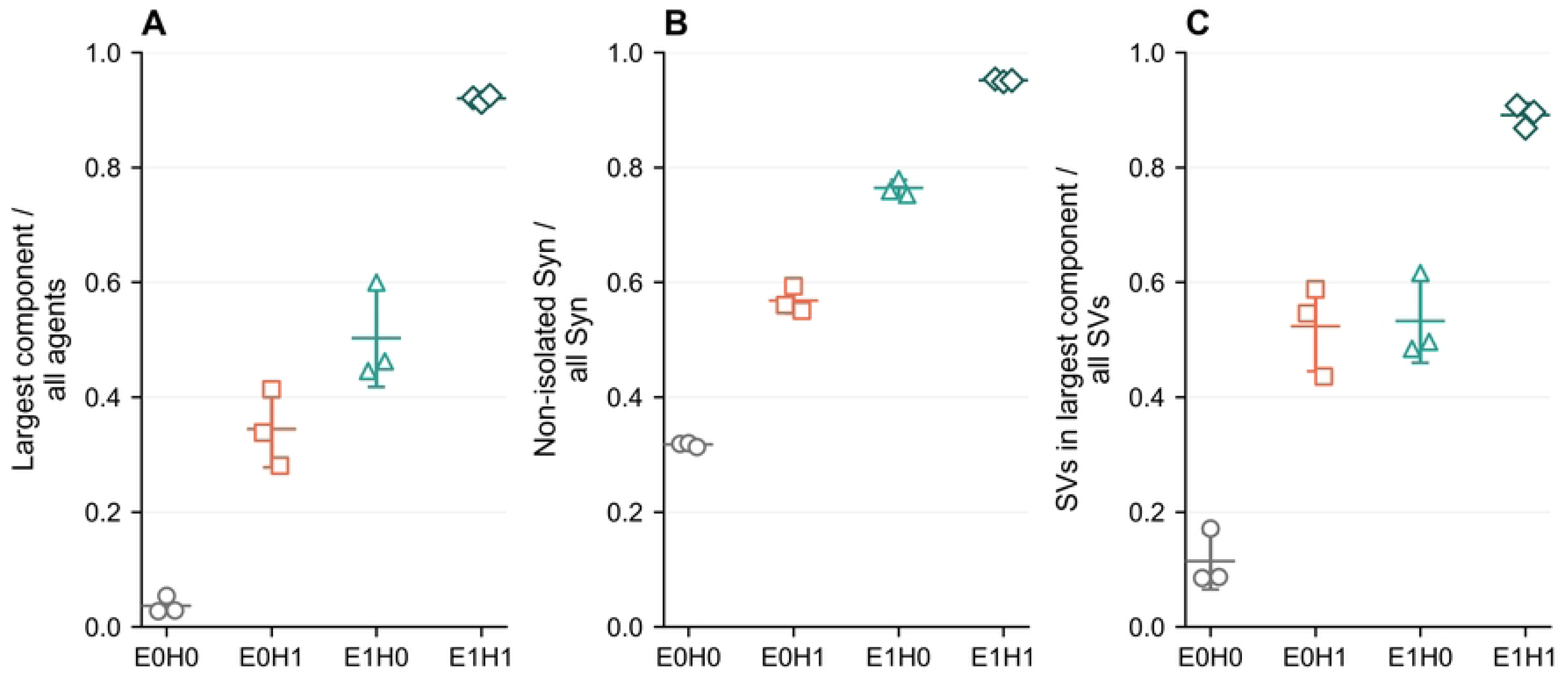
Structural outcomes in the final observation window. The graph shell is 2 nm and the averaging window is 58.94–84 ms. (A) Fraction of all agents in the largest component. (B) Fraction of Syn agents in components of two or more agents. (C) Fraction of vesicles in the largest component. Open symbols show individual runs. Horizontal marks and bars show means and one between-seed SD (*n* = 3 per condition). E and H indicate electrostatic and hydrophobic interactions (1 active, 0 inactive). Van der Waals attraction is active throughout.

**Table 1.** Structural outcomes over 58.94–84 ms at the primary 2-nm association shell.

| Condition | All-agent largest component | Non-isolated Syn | SVs in largest component |
|---|---|---|---|
| E0H0 | 0.0368 (0.0150) | 0.3172 (0.0035) | 0.1143 (0.0492) |
| E0H1 | 0.3440 (0.0667) | 0.5678 (0.0221) | 0.5229 (0.0783) |
| E1H0 | 0.5025 (0.0847) | 0.7639 (0.0142) | 0.5323 (0.0728) |
| E1H1 | 0.9200 (0.0064) | 0.9516 (0.0023) | 0.8909 (0.0204) |

Entries are means across three seeds with sample standard deviations (SDs) in parentheses. Van der Waals attraction is present in all conditions.

The assemblies developed at different rates (Fig 2A). At 28 ms, the mean all-agent largest- component fractions were 0.020, 0.090, 0.099 and 0.610 for E0H0, E0H1, E1H0 and E1H1. At 84 ms, they were 0.034, 0.409, 0.595 and 0.926. Growth in E1H1 slowed towards the end. The mean fitted slope decreased from 0.0148 per ms over 14–28 ms to 0.00038 per ms over 70–84 ms (Fig 2C). The single-interaction conditions continued to grow during the extended observation.

**Fig 2.**
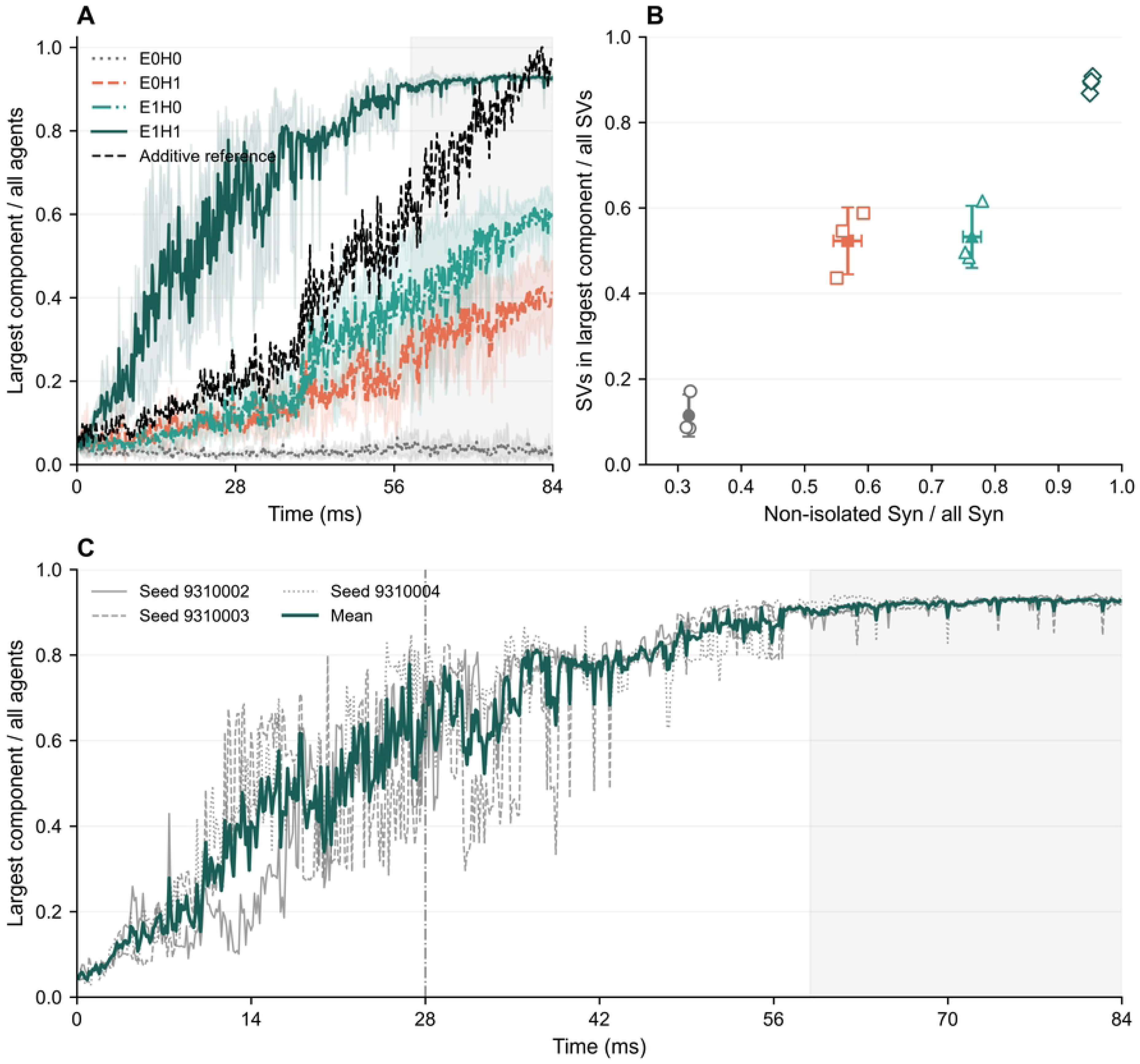
Network growth and vesicle participation. The graph shell is 2 nm. (A) All-agent largest- component fraction for all four conditions through 84 ms. Lines show condition means, bands show one between-seed SD, and the dashed line is the additive reference *L*_10_(*t*) + *L*_01_(*t*) − *L*_00_(*t*). Shading marks the final 180 frames. (B) Late-window non-isolated Syn fraction versus SV participation in the largest component. Open symbols show runs. Filled symbols and bars show means and one SD. (C) The three E1H1 trajectories and their mean. The vertical line marks 28 ms and shading marks 58.94–84 ms.

For each seed, we compared joint activation with the sum of the separate E and H effects relative to E0H0. The interaction contrast is

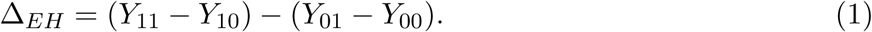

For the all-agent largest-component fraction, the mean contrast over 19.74–28 ms was 0.3723, with positive values in all three seeds. Over 58.94–84 ms, its mean decreased to 0.1103, with seed values 0.1660, −0.0463 and 0.2112. Growth under E or H alone reduced the initial separation from their additive reference.

### 2.2 Protein association and vesicle participation follow different time courses

Electrostatics produced the larger increase in the non-isolated Syn fraction *Q*_Syn_. At 84 ms, its late-window mean was 0.7639 with E alone and 0.5678 with H alone (Table 1). The corresponding SV fractions in the largest component, *L*_SV_, were similar: 0.5323 and 0.5229. The two conditions therefore had different protein participation despite comparable collective vesicle participation.

This relationship developed over time. In the earlier 19.74–28-ms window, the non-isolated Syn fraction was higher with E alone than with H alone (0.6989 versus 0.5157), but fewer SVs in the largest component (0.1273 versus 0.1839). That ordering of vesicle participation held in every seed at 28 ms. The longer observation shows that it describes an early stage of assembly.

With both interactions active, the 84-ms late-window means reached 0.9516 for *Q*_Syn_ and 0.8909 for *L*_SV_. Their E-by-H contrasts were −0.0628 and −0.0500, respectively. For *L*_SV_, the seed contrasts were −0.0369, −0.1642 and 0.0509, compared with three positive values and a mean of 0.3811 at 28 ms. The conditional differences are given in Table 2. The interaction contrast thus depends on the structural observable and the observation period.

**Table 2.** Paired seed-block contrasts over 58.94–84 ms at the 2-nm shell.

| Outcome | Conditional contrast | Mean difference (SD) |
|---|---|---|
| All-agent largest component | H effect at E0 | 0.3072 (0.0538) |
|  | H effect at E1 | 0.4175 (0.0906) |
|  | E effect at H0 | 0.4656 (0.0698) |
|  | E effect at H1 | 0.5760 (0.0730) |
| Non-isolated Syn | H effect at E0 | 0.2506 (0.0194) |
|  | H effect at E1 | 0.1877 (0.0159) |
|  | E effect at H0 | 0.4466 (0.0115) |
|  | E effect at H1 | 0.3838 (0.0238) |
| SVs in largest component | H effect at E0 | 0.4086 (0.0562) |
|  | H effect at E1 | 0.3586 (0.0929) |
|  | E effect at H0 | 0.4180 (0.0239) |
|  | E effect at H1 | 0.3680 (0.0900) |

Values are mean differences in fractions, with the SD across the three paired differences in parentheses. The H effect compares H1 with H0 at fixed E. The E effect compares E1 with E0 at fixed H.

### 2.3 Local vesicle coverage and connections between groups

We separated Syn–Syn, Syn–SV and SV–SV contacts in the saved frames. At the 2-nm shell, the 84-ms late-window fraction of Syn agents next to at least one SV was 0.1906 in E0H0, 0.4647 in E0H1, 0.7508 in E1H0 and 0.9499 in E1H1 (Fig 3).

**Fig 3.**
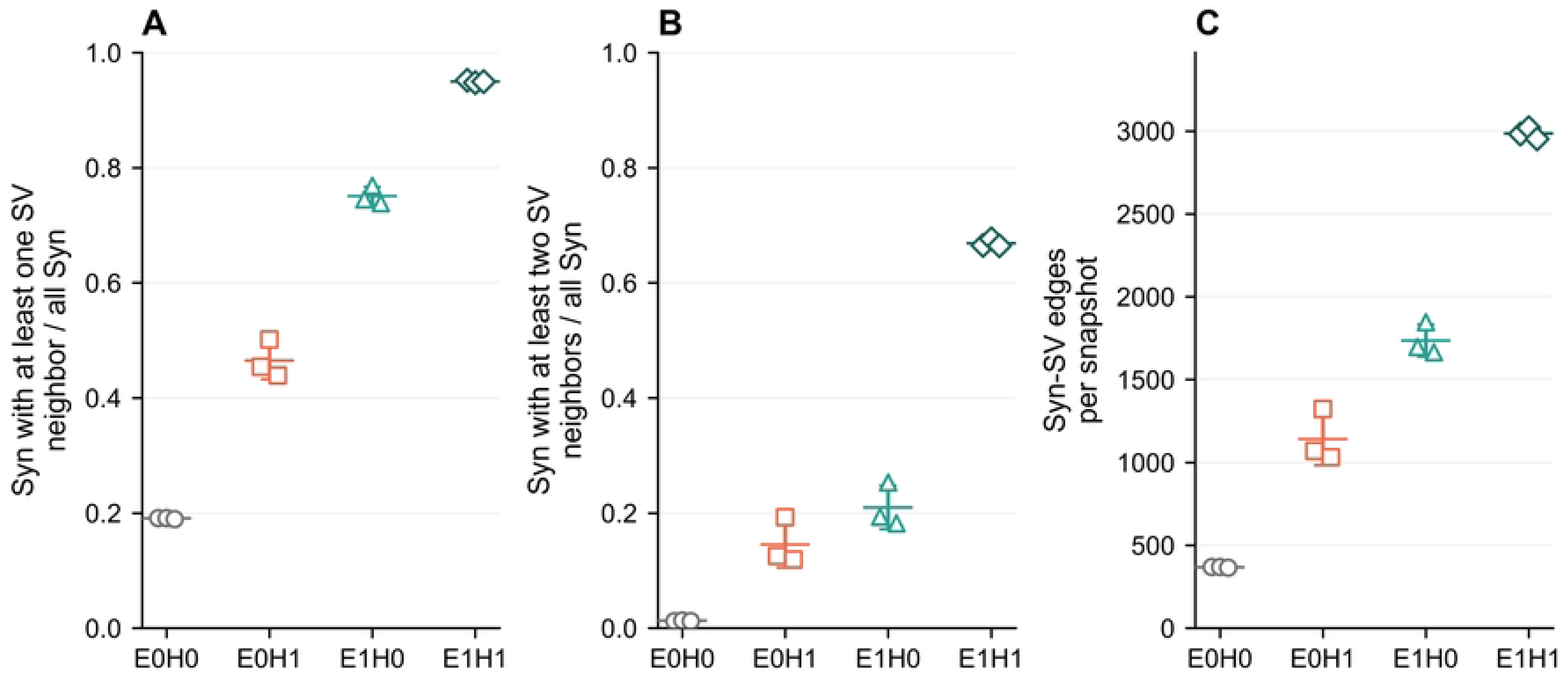
Species-resolved contacts. Values are averaged over 58.94–84 ms at the 2-nm graph shell. (A) Fraction of Syn agents adjacent to at least one SV. (B) Fraction adjacent to at least two distinct SVs. (C) Number of Syn–SV edges per snapshot. Open symbols show individual late-window run means. Marks and bars show the mean and one between-seed SD (*n* = 3 per condition). Contact uses outer radii, including the SV corona.

Over 19.74–28 ms with electrostatics alone, 99.72% of SVs had a Syn neighbor, while 12.73% belonged to the largest component. At 84 ms, the corresponding late-window values were 99.14% and 53.23%. Extensive local coverage therefore preceded collective vesicle connectivity. In E1H1, 98.08% of SVs had a Syn neighbor and 89.09% belonged to the largest component in the later window.

The late-window fraction of Syn agents next to at least two SVs was 0.1454 with H alone, 0.2097 with E alone and 0.6688 with both interactions (Fig 3B). Mean Syn–SV edge counts per snapshot were 1,140.6, 1,735.7 and 2,985.4, respectively. The joint condition also had more Syn–Syn edges, with a mean of 1,372.8, compared with 469.2 under H alone and 314.5 under E alone. Its many multivesicle neighbors provide geometric connections between vesicles within the association graph. The single-interaction conditions differed in contact composition. E1H0 had more total edges than E0H1 (2,082.0 versus 1,793.9 per snapshot), but fewer Syn–Syn and SV–SV edges. Their similar *L*_SV_ values at 84 ms corresponded to different local contact networks. Their all-agent largest-component fractions remained more distinct because that measure also counts Syn agents.

### 2.4 The joint condition retains the largest assembly across observation shells

Across 0, 2, 4 and 6 nm surface-gap shells, E1H1 retained the largest mean all-agent component and largest-component SV fraction at 84 ms. Its all-agent fraction ranged from 0.8872 to 0.9382, compared with 0.0185–0.1433 for E0H0. The ordering was consistent across these graph thresholds applied to the same trajectories.

### 2.5 Stronger assembly accompanies slower short-lag motion

We estimated Syn mobility from saved positions projected onto the *xy* plane. Apparent diffusion coefficients came from fits to each agent’s time-averaged mean-squared displacement (MSD) over lags of 2.10–3.92 ms. For the full 84-ms trajectories, across-seed means of the agent-mean apparent coefficient were 0.1403, 0.1308, 0.1108 and 0.0626 µm^2^ s^−1^ for E0H0, E0H1, E1H0 and E1H1, respectively (Fig 4). Mobility was about 55% lower in E1H1 than in E0H0.

**Fig 4.**
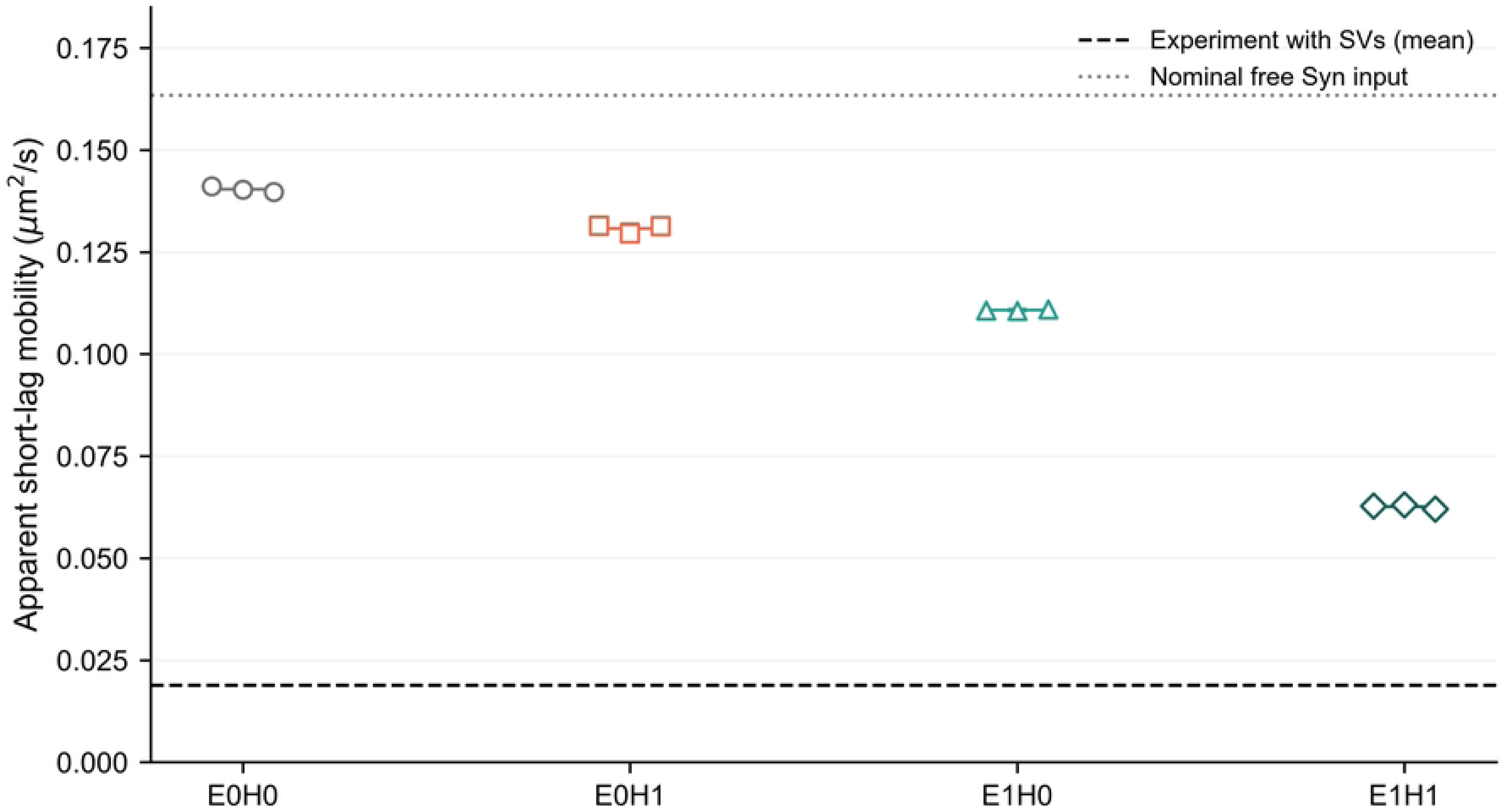
Short-lag synapsin mobility and its experimental reference. Simulation estimates use the full 84-ms trajectories. Symbols show run means over Syn agents. Marks and bars show the between-seed mean and one SD. The apparent coefficient is one quarter of the free-intercept slope of projected time-averaged MSD over 2.10–3.92 ms. The dashed line is the Hoffmann et al. with-SV mean from released Fig. 2h source data [8]. The dotted line shows the nominal unconfined Syn diffusion coefficient.

The source data of Hoffmann et al. give a mean short-time synapsin coefficient of 0.0189 µm^2^ s^−1^ with SVs, versus 0.0253 µm^2^ s^−1^ without SVs [8]. The E1H1 full-trajectory mean is about 3.3 times the experimental with-SV mean. Restricting the simulation analysis to 58.94–84 ms gives an E1H1 mean of 0.0548 µm^2^ s^−1^. S1 Text gives the per-run estimates for both observation windows.

## 3 Discussion

### 3.1 From local association to mesoscale vesicle connectivity

Electrostatics brought Syn agents close to almost every vesicle early, while most vesicles remained in separate groups. With hydrophobic attraction alone, more vesicles participated in the largest component over the earlier window despite lower Syn participation. By the later window, vesicle participation was similar under the two interactions alone, although their contact networks still differed.

The signs of the forces help explain these networks (Table 3). Electrostatics attracts Syn–SV pairs and repels like-species pairs. Hydrophobic attraction acts on Syn–Syn and Syn–SV pairs. Consistent with these differences, electrostatics alone favored Syn–SV contacts, while hydrophobic attraction alone yielded more Syn–Syn and SV–SV contacts. With both interactions active, many more Syn agents were close to several vesicles. Each such contact provides a path between vesicles in the association graph.

**Table 3.** Pair-level action of the two interaction terms at positive outer-surface gap.

| Pair class | Electrostatic interaction when E1 | Hydrophobic interaction when H1 |
|---|---|---|
| Syn–Syn | Repulsive | Attractive |
| Syn–SV | Attractive | Attractive |
| SV–SV | Repulsive | Zero amplitude |

Signs describe radial forces for the assigned effective surface potentials. Dispersion attraction acts in all three pair classes and remains enabled in every condition.

Connectivity changes nonlinearly with the number and placement of edges, as in random graphs and continuum percolation [20, 21]. Here, the large positive contrast in the all-agent component decreased as the single-interaction assemblies grew. By 84 ms, its sign varied across seeds, and the mean vesicle-participation contrast was slightly negative. The joint condition still had the largest assembly. The strength of the early cooperative response therefore depends on the time window and the observable.

### 3.2 Connectivity as a distinct feature of vesicle clusters

Protein enrichment in a condensate and connectivity across a vesicle cluster describe different properties. The simulations show how broad local association can precede the growth of a connected assembly. We propose that LLPS could support vesicle clustering by creating conditions in which local protein–vesicle contacts link separate groups. Measuring synapsin partitioning and vesicle connectivity together would help test that interpretation during perturbations of protein–membrane and protein–protein interactions.

### 3.3 Relation to molecular and particle-level models

The particle model of Alfken et al. linked redistribution of vesicle-bound synapsin to adhesion and finite assembly size [13]. Their two-dimensional periodic Langevin model follows proteins on vesicle surfaces. Our model follows freely moving Syn agents inside a three-dimensional compartment and separates electrostatic from hydrophobic interactions.

Residue-level simulations describe charge and hydropathy along protein chains [17]. In repetitive peptides, hydrophobic attraction competes with same-sign electrostatic repulsion [18]. Our two-species model represents this competition for Syn–Syn pairs and includes attraction between oppositely charged Syn–SV pairs. The latter favors local protein association, while larger vesicle assemblies also depend on how the contacts connect separate groups.

Atomistic simulations and experiments connect an ATP-dependent synapsin conformational transition to vesicle binding and oligomerization [16]. At a coarser resolution, Moussavi-Baygi et al. used Brownian dynamics with effective hydrophobic interactions to investigate selective transport through the nuclear pore complex [22]. Jamali et al. also used an agent-based model to examine how ligand organization, receptor self-association and binding affinity contribute to integrin clustering [23]. These studies provide examples of how effective local interactions can be used to study transport or clustering at larger scales.

### 3.4 Experimental context and biological implications

Scattering measurements show noncompact synapsin–vesicle clusters [14]. Connected-component measures can complement measurements of cluster size in such structures. In non-neuronal cells, Park et al. found that synapsin and synaptophysin together organized SV-like vesicles into liquid- like clusters [9]. Kim et al. identified an electrostatic contribution to coacervation of the two proteins [24]. Yoshida et al. found that endophilin A1 promoted synapsin condensation and entered SV-like clusters in a synapsin-dependent manner. Their correlative microscopy also revealed endophilin–synapsin condensates without vesicle membranes [10]. This observation distinguishes protein condensation from the incorporation of vesicles.

The distinction between coverage and connectivity may also help interpret synapsin dysfunction. Human *SYN1* loss-of-function variants associated with autism and epilepsy impair vesicle-pool size and trafficking [25]. Oxidation and persulfidation have been linked to different changes in synapsin condensation and neurotransmission [26]. Together, these studies motivate experiments that measure local vesicle association and network connectivity.

### 3.5 Scope and limitations

The simulations cover one concentration and compartment volume. At 84 ms, growth in E1H1 had slowed and the single-interaction assemblies were still developing. The contrasts apply to the observation windows reported here, and their precision is limited by the three seed blocks. Changing concentration could shift the response near a connectivity threshold or saturation.

The model represents synapsin and vesicles as spheres with effective surface forces. Each switch changes more than one pair class, and geometric contacts measure proximity. Native terminals also contain endocytic proteins that redistribute during activity [27] and active-zone scaffolds that organize docking and release [28]. The model omits these components along with explicit protein valency and phosphorylation. It also omits membrane deformation, active vesicle cycling and hydrodynamic coupling. The force switches are effective perturbations of the model.

The numerical checks establish geometric admissibility. Convergence of the reported observables at smaller integration steps remains to be assessed. Establishing LLPS would require additional evidence on phase coexistence, molecular exchange and system-size dependence.

The mobility comparison uses condensate-core tracks sampled at 1 ms in Hoffmann et al. and all Syn agents sampled at 0.14 ms. Simulated tracks have no localization error or exposure blur, and each simulated condition contains vesicles. Effective friction was chosen with reference to published mobility scales. The experimental mean provides a scale reference for the apparent mobility. Quantitative comparison would require matched tracking conditions.

## Conclusion

In the model, electrostatics promotes early coverage of vesicles by Syn agents. With hydrophobic attraction also active, a larger connected assembly forms sooner and Syn mobility is lower. The contrast between the joint response and the sum of the separate responses changes as the single- interaction assemblies grow. Tracking local contacts and network connectivity separately captures this evolution and provides a way to examine vesicle organization alongside protein enrichment.

## 4 Materials and methods

### 4.1 Simulation design and numerical settings

The four force conditions each have three independent seed blocks (9310002–9310004), giving 12 trajectories. Every run contains 200 SVs and 1,807 Syn agents in a closed 0.1 µm^3^ compartment. Each trajectory comprises 120,000 integration steps and 600 saved frames. The original 28-ms trajectories were continued from checkpoints after inspection showed continuing growth. Physical parameters and the random number generator state were retained. Duplicate join frames were counted once. The numerical settings are common to all conditions. S1 Text gives the per-run numerical diagnostics.

### 4.2 Compartment, size distributions, and population

We represent SVs and synapsin as spherical agents in three dimensions, following the scale of synapsin–vesicle reconstitution and neuronal imaging [6, 8]. The closed spherical compartment has volume *V* = 0.10 µm^3^, corresponding to radius

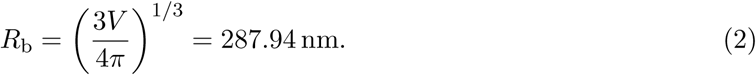

The scale is informed by three-dimensional hippocampal bouton reconstructions [29]. An SV has membrane-core radius *b_i_*, corona thickness *s_i_* = 4.8 nm, and outer radius *a_i_* = *b_i_* + *s_i_*. Each Syn agent has *b_i_* = *a_i_* = 3.972 nm and *s_i_* = 0. The corona represents the protein-rich vesicle surface [14]. In the model, coronas can interpenetrate.

SV core radii are independently rejection-sampled from a normal distribution with location *µ_R_* = 20.45 nm and input SD *σ_R_* = 2.5 nm, restricted to 15 ≤ *b_i_* ≤ 30 nm. The location and SD inputs follow the excitatory-SV diameter measurements of Tao et al. [30]. The bounds are informed by cryo-electron microscopy (cryo-EM) and small-angle X-ray scattering (SAXS) measurements [31]. These values specify the distribution before truncation. The number of Syn agents is calculated

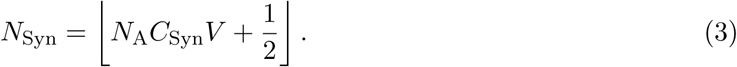

Here *N*_A_ is Avogadro’s constant, with *C*_Syn_ in mol*/*m^3^ and *V* in m^3^. The floor expression ⌊*x* + 1*/*2⌋ rounds a nonnegative value to the nearest integer. The chosen 30 µmol*/*L concentration gives *N*_Syn_ = 1807. There are 200 SVs at the canonical volume. Initial centers are sampled uniformly within active cells and restricted to the particle-accessible sphere. Before the physical trajectory, an overlap-relaxation stage reduces initial core intersections. Its stopping criterion is an overlap below 0.05 nm for three checks or no overlapping pairs. The stage stops after at most 200 iterations.

### 4.3 Linked cells and boundary treatment

The supremum outer SV diameter is *d*_max_ = 2(30 + 4.8) = 69.6 nm. For each bounding-box dimension *L_d_*, the grid uses

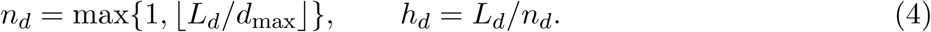

The canonical sphere has eight cells per bounding-box axis, with *h_d_* = 71.9853 nm. Cells are assigned by particle center. The force calculation searches the home cell and two neighboring cell layers. The linked-cell stencil defines the full set of candidate pairs for the force calculation.

The largest center distance at a 6-nm outer-surface gap is *d*_max_ + 6 = 75.6 nm, within the two-layer search distance of 143.9706 nm. A particle center is confined to radius *R*_b_ − *a_i_* from the compartment center. Out-of-domain proposals are mapped inward by radial folding. The same

map is applied after overlap corrections. The resulting boundary approximates a closed reflecting wall.

### 4.4 Pair interactions

Let *r_ij_* = ||***x****_i_* − ***x****_j_*||, ***n****_ij_* = (***x****_i_* − ***x****_j_*)*/r_ij_*, and *g_ij_* = *r_ij_* − *a_i_* − *a_j_*. A positive radial force is repulsive. The production force definition is

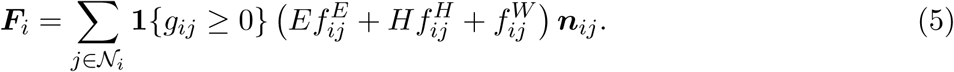

Here N*_i_* is the linked-cell candidate set and *E, H* ∈ {0, 1} switch the electrostatic and hydrophobic terms. Van der Waals attraction remains active in every condition. At negative outer gap, the surface-force terms vanish and geometric correction controls core overlap.

The electrostatic kernel is a reduced-potential, screened sphere interaction:

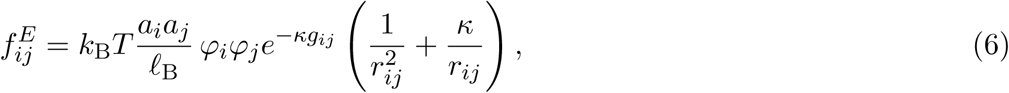

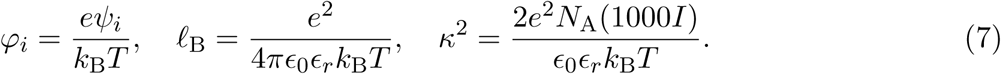

The last expression uses SI quantities and *I* in mol*/*L. Lengths are then converted to nanometers.

At *T* = 310.15 K, *ɛ_r_* = 74.2, and *I* = 0.15 mol*/*L, κ^−1^ = 0.77886 nm. SV and Syn effective potentials are respectively −22.5 and +16 mV. Negative isolated-SV zeta potentials motivate the SV input [32]. The Syn input represents an effective positive protein charge scale. The kernel approximates screened electrostatics in a linearized-Poisson–Boltzmann/Yukawa form.

Hydrophobic attraction is

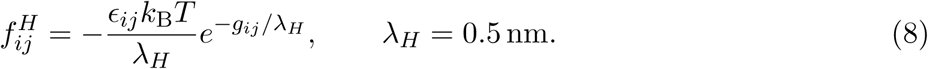

This follows from 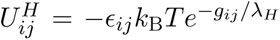 outside contact. An exponential kernel has an experi- mental precedent in surface-force measurements [33]. Its decay length and pair amplitudes specify the effective short-range attraction. The SV–SV hydrophobic amplitude is zero.

For the fixed dispersion background, the membrane/core radii are used:

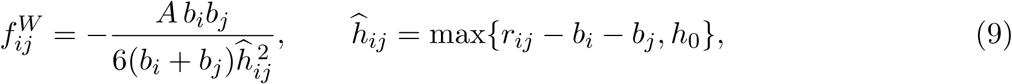

where *A* = 5 × 10^−21^ J and *h*_0_ = 0.3 nm. This non-retarded Hamaker–Derjaguin approximation is informed by dispersion theory [34]. We use a common effective constant for the aqueous back- ground. The dispersion surfaces are defined by core radii, with lower-gap regularization to bound the attraction near contact.

### 4.5 Mobility and discrete stochastic dynamics

Independent-particle Stokes drag gives

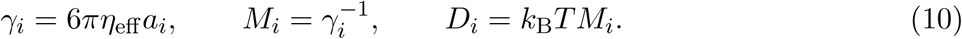

Brownian particle updates use diagonal mobility [35]. The effective friction input is *η*_eff_ = 0.35 Pa s, selected with reference to published mobility scales [8, 36]. The nominal unconfined diffusion coefficients are 0.1634 µm^2^ s^−1^ for a Syn agent and 0.02571 µm^2^ s^−1^ for an SV with *a_i_* = 25.25 nm.

At each Euler–Maruyama step,

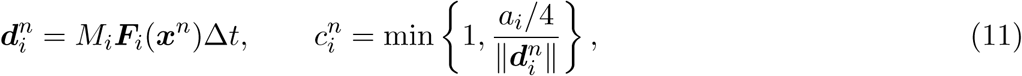

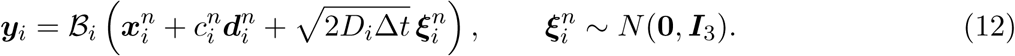

For zero drift, *c^n^* = 1. The limiter acts on deterministic displacement, while the Brownian increment remains Gaussian. B*_i_* is the boundary map. The next state is the result of mobility-weighted iterative overlap correction applied to ***y***. Pair admissibility is

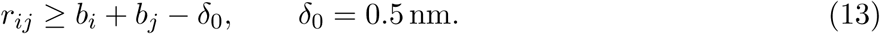

Thus permitted outer overlaps are 0.5 nm for Syn–Syn, 5.3 nm for Syn–SV, and 10.1 nm for SV– SV. Corrections are accumulated simultaneously, averaged over each particle’s violating contacts, reflected at the wall, and repeated with updated cells. The residual tolerance is 0.005 nm, with early stopping and a 128-iteration safety ceiling.

The fixed integration step is Δ*t* = 0.0007 ms. Each trajectory has 120,000 physical moves over 84 ms. Snapshots are scheduled every 200 steps, or 0.14 ms. Forces are evaluated at the stored state before movement, aligning saved forces and positions.

### 4.6 Structural graphs and observables

A nonperiodic undirected graph is constructed independently at each stored frame using *A*^(^*^ℓ^*^)^ = **1**{*g_ij_* ≤ *ℓ*}, *i* = *j*. The primary outer-surface shell is *ℓ* = 2 nm, with 0, 2, 4, 6 nm sensitivity shells. Edges represent geometric association. Let C*_f_* be the connected components at frame *f*, *n_c_* = |*c*|, and *c^∗^* the component with the largest total agent count. The species-resolved structural observables are

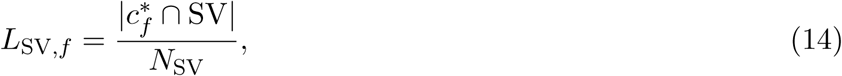

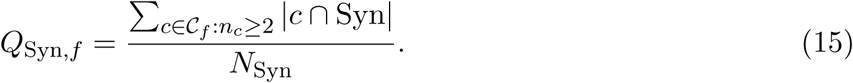

Thus *L*_SV_*_,f_* counts vesicles in the component selected by total agent count, and *Q*_Syn_*_,f_* counts Syn agents in any component containing at least two agents, including Syn–Syn groups. The primary structural observable is the all-agent largest component fraction

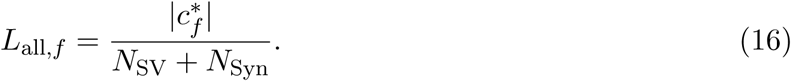

Each outcome is averaged over the last *m* = max{1, ⌈0.30*F* ⌉} of the *F* stored frames:

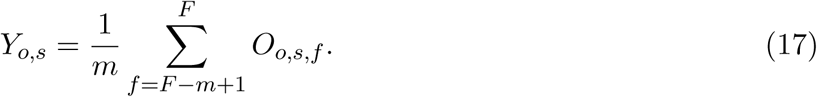

The indices *o*, *s* and *f* denote the observable, seed and saved frame. We suppress the force-condition index. Each run is summarized before values are compared across seeds. For these trajectories, the 84-ms late window covers frames 421–600 (58.94–84 ms). The earlier comparison uses the same 30% rule on the first 200 frames, giving frames 141–200 (19.74–28 ms).

### 4.7 Force-factorial design

The factorial design fixes ionic strength at *I* = 0.15 mol*/*L and nominal synapsin concentration at 30 µmol*/*L. It includes all four combinations of the binary interaction switches, *E, H* ∈ {0, 1}. H1 uses the reference amplitudes *ɛ*_SynSyn_ = 1.50 and *ɛ*_SynSV_ = 1.00, corresponding to well depths of 1.50*k*_B_*T* and 1.00*k*_B_*T*, respectively, with *ɛ*_SVSV_ = 0. H0 sets all hydrophobic amplitudes to zero. E1 uses the electrostatic kernel of eq. (6) and E0 sets that kernel to zero. In every condition, the same van der Waals law and its parameters remain active.

Three seed blocks, 9310002–9310004, were completed in all four conditions. An earlier four-run pilot with seed 9310001 informed feasibility and the selection of structural observables. The 12 later trajectories provide the comparisons reported here. The 2-nm primary shell and the three sensitivity shells were selected before those runs. Fixed inputs are listed in Table 4.

**Table 4.** Fixed inputs of the four-condition force comparison.

| Quantity | Value | Model role |
|---|---|---|
| Integration step | 0.0007 ms | Fixed in every condition |
| Duration | 84 ms | 120,000 physical steps |
| Saved-frame spacing | 0.14 ms | 600 saved frames per completed run |
| Temperature | 310.15 K | Shared setting |
| Effective viscosity | 0.35 Pa s | Literature-informed effective friction |
| Relative permittivity | 74.2 | Solvent approximation |
| Volume | 0.10 µm<sup>3</sup> | Closed spherical compartment |
| Synapsin concentration / count | 30 *µ*M / 1,807 | Same in all conditions |
| Vesicle count | 200 | Same in all conditions |
| Vesicle core-radius input | 20.45 nm | Bounds: 15–30 nm |
| Core-radius SD input | 2.5 nm | Truncated heterogeneous input |
| Corona thickness | 4.8 nm | Effective interpenetrable shell |
| Syn effective radius | 3.972 nm | Coarse-grained input |
| SV / Syn surface potentials | −22.5 / +16 mV | Effective surface-potential inputs |
| Hamaker constant | 5 × 10<sup>−21</sup> J | Fixed van der Waals background |
| Surface regularization | 0.3 nm | Numerical/force definition |
| Hydrophobic decay length | 0.5 nm | Effective short-range input |
| H1 well depths Syn–Syn / Syn–SV | 1.50 / 1.00 *k*<sub>B</sub>*T* | Syn–Syn and Syn–SV only. H0 sets zero |
| Core overlap / residual | 0.5 / 0.005 nm | Distinct geometric quantities |
| Projection iteration cap | 128 | Early stopping enabled |
| Neighbor depth | Two cell layers | Candidate-pair force evaluation |
| Primary analysis shell | 2 nm | Sensitivity: 0, 2, 4, 6 nm |

The binary E and H switches are the only inputs that vary by condition.

### 4.8 Post hoc mobility analysis and experimental reference

Fig. 2h of Hoffmann et al. summarizes apparent diffusion coefficients from synapsin trajectories projected onto the imaging plane [8]. The coefficients were evaluated over 2–4 ms with 1000-Hz tracking. We use the mean with SVs from their source data as an experimental reference.

To match the dimensionality of those measurements, we project the Syn trajectories onto the *xy* plane. We convert coordinates from nm to µm and write ***z****_i,f_* = (*x_i,f_, y_i,f_*) for agent *i*. With *F* = 600 and *δt*_save_ = 0.14 ms, the time-averaged squared displacement at lag *k* is

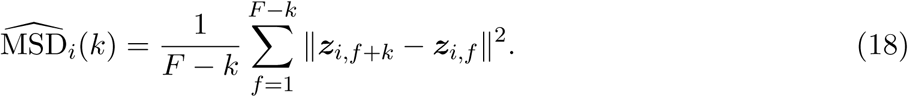

We fit M^^^SD*_i_*(*k*) to *α_i_* + *β_i_τ_k_* by ordinary least squares with a free intercept, using all 14 native lags from 2.10 to 3.92 ms. The lag time *τ_k_* = *k δt*_save_ is expressed in seconds. We define the apparent diffusion coefficient as *D*^^^*i* = *β_i_/*4, in µm^2^ s^−1^. The run summary is the mean of *D*^^^*i* over 1,807 Syn agents. We then report the mean and SD across the three runs in each condition. The estimator is also applied to the final 180 frames to assess window sensitivity. S1 Text gives the per-run estimates and fitted-slope diagnostics. S1 Data includes the agent-level values.

### 4.9 Post hoc species-resolved contact analysis

For every saved frame, the same 2-nm association graph is reconstructed from recorded centers and outer radii. Each unordered edge is counted once and classified as Syn–Syn, Syn–SV or SV–SV.

Let S and V be the Syn and SV index sets. For Syn agent *i* ∈ S, let

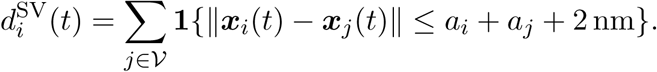

The fractions with at least one or at least two SV neighbors are 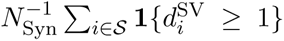and 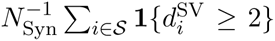. The reciprocal SV coverage counts SVs with at least one Syn neighbor, divided by *N*_SV_. Counts and fractions are first averaged within each run over its last 180 frames, and only then summarized across seeds. The analysis uses the same saved trajectories and 2-nm shell as the structural comparison.

### 4.10 Seed-level statistical analysis

Each complete trajectory is one replicate, with frames, agents and clusters nested within it [37]. We pair the run-level comparisons by seed across conditions.

Each structural observable is first averaged over the final 180 frames of a trajectory. We report the mean and sample SD of the three run-level values for each condition. For a comparison of conditions *A* and *B*, the paired difference is *d_s_* = *Y_B,s_* − *Y_A,s_*. We summarize the three differences by their mean and sample SD. The E-by-H interaction is calculated within each seed block using eq. (1). Individual seed values are shown in the figures and S1 Text. We use the size and consistency of paired differences to interpret the results.

### 4.11 Use of AI

The authors used ChatGPT and Codex (OpenAI) to support code development and refine the language of the manuscript. Claude (Anthropic) provided feedback on early drafts. The authors remain responsible for checking the code, mathematical definitions, primary sources, numerical results, and interpretation, and for approving the submitted manuscript.

## Author contributions

Shokoofeh Akbari: Conceptualization, Methodology (mathematical formulation), Software, Investigation, Formal analysis, Data curation, Visualization, and Writing–original draft.

Oleg Shupliakov: Conceptualization (biological and LLPS input) and Writing–review and editing. Yousef Jamali: Conceptualization, Methodology (scientific interpretation), Supervision, Funding acquisition, and Writing–review and editing.

## Data and code availability

The numerical data underlying the figures and tables are provided in the Supporting Information. The reproducibility package contains the simulator and the 12 condition configurations, together with analysis scripts, processed per-seed results, numerical diagnostics and trajectory checksums. It is maintained at https://github.com/BioMathLab-TMU/simulationsv-electrostatic-hydrophobic-response, version manuscript-84ms-20261030, which contains the common 84-ms inputs and reproduction scripts. Release v1.0.0 contains the original 28-ms inputs. Original code is licensed under MIT. Author-generated data and tables are licensed under CC BY 4.0. Third- party code and experimental data retain their original licenses. The experimental source data of Hoffmann et al. are available at https://doi.org/10.6084/m9.figshare.24179445.v1.

## Supporting information captions

**S1 Text. Supporting methods and numerical results.** Per-seed structural endpoints, paired primary-interaction estimates, association-shell sensitivity, interface definitions and complete run- level interface outcomes, together with statistical and software validation details. (PDF)

**S1 Data. Numerical data underlying the figures and tables.** Per-seed structural observables, paired contrasts, graph-shell sensitivity, time-series inputs, interface diagnostics and mobility comparison values. (ZIP)

